# A ternary switch determines ERα LBD conformation

**DOI:** 10.1101/2025.02.09.637324

**Authors:** Daniel P. McDougal, Jordan L. Pederick, Scott Novick, Blagojce Jovcevski, Annmaree K. Warrender, Bruce D. Pascal, Patrick R. Griffin, John B. Bruning

**Affiliations:** Institute for Photonics and Advanced Sensing (IPAS), School of Biological Sciences, The University of Adelaide, Adelaide, SA 5005, Australia; Department of Molecular Medicine, The Herbert Wertheim UF Scripps Institute for Biomedical Innovation & Technology, Jupiter, FL 33458; Department of Molecular and Biomedical Sciences, School of Biological Science, The University of Adelaide, Adelaide, SA 5005, Australia; Australian Synchrotron, Australian Nuclear Science & Technology Organisation, 800 Blackburn Road, Clayton VIC, 3168, Australia; Omics Informatics LLC. 1050 Bishop Street #517, Honolulu, HI 96813

## Abstract

The transcription factor estrogen receptor α (ERα) is the primary driver of ER+ breast cancer progression and a target of multiple FDA-approved anticancer drugs. Ligand-dependent activity of ERα is determined by the conformation of helix-12 (H12) within the ligand binding domain (LBD), but how H12 transitions from an unliganded (apo) state to active (estrogen-bound) or inactive (SERM/SERD-bound) states remains unresolved. Here, we present the first crystal structure of an apo ERα LBD, revealing a third distinct H12 conformation that regulates receptor activity. Structural mass-spectrometry, small-angle X-ray scattering, functional analysis and molecular dynamics simulations reveal that the apo conformation of H12 is stable in the absence of ligand, but is destabilised by Y537S and D538G breast cancer mutations driving constitutive activation. We propose a model in which H12 functions as a ternary molecular switch to determine receptor activity. These findings provide critical insights into the ligand-dependent and -independent regulation of ERα and have significant implications for therapeutic intervention.

## Introduction

Estrogen receptor α (ERα) is a ligand-responsive nuclear hormone receptor transcription factor that regulates physiological and cellular processes across vertebrates including reproduction, cellular proliferation and survival (*1*). In line with this, ERα signalling has been identified as a key driver of breast cancer development and progression, of which at least 70% are ER+ (*2, 3*). Typically, binding of ligands to the ERα LBD determines the transcriptional outcome of the receptor through stabilizing discrete conformational states of the C-terminal helix, H12 (**Fig. 1A**) (*4*). The central model for ligand-dependent activation of nuclear receptors is that ligand binding switches H12 from a dynamic state to one that is stable (*5, 6*). However, there is evidence that this is not the case for the ERα LBD. Hydrogen-deuterium exchange coupled mass-spectrometry (HDX-MS) analysis of the human ERα LBD shows that ligand binding shifts protein structure from a dynamic ensemble toward a stabilised state. Yet, no significant differences in the deuterium uptake of H12 occurs, supporting that H12 exists in a stable, discrete conformation prior to ligand binding (*7–10*).

**Figure 1.**
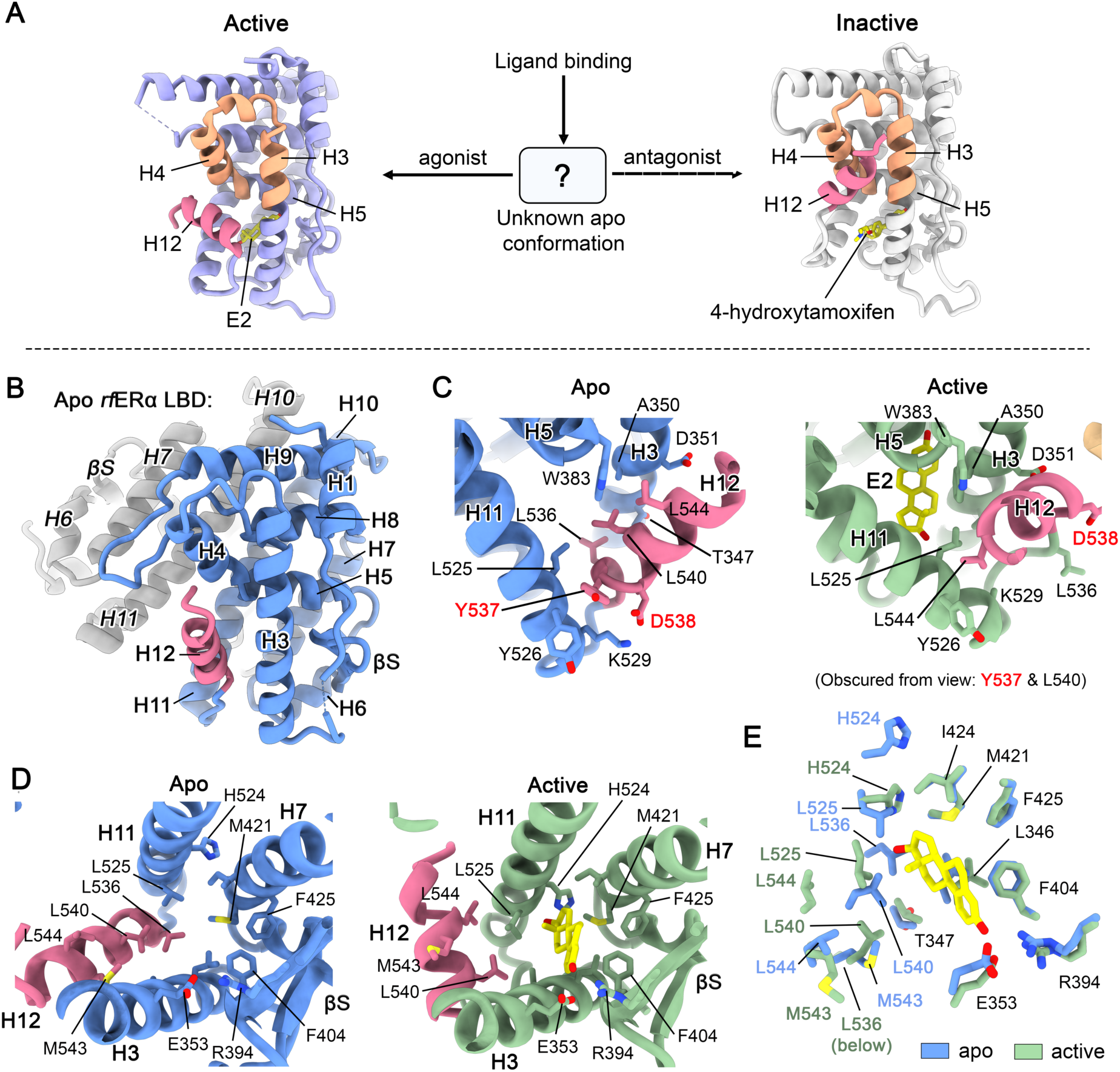
Structural comparison of the apo and E2-bound states of the rfERα LBD. **A**) Cartoon schematic illustrating that the conformation of H12 prior to ligand binding is unknown. Left: *h*ERα LBD bound to E2 (PDB: 3UUD); right: *h*ERα LBD bound to the SERM 4-hydroxytamoxifen (PDB: 3ERT). In the schematic, H12 is coloured pink while helices 3, 4, and 5, which form the AF2, are coloured orange. E2 and 4-hydroxytamoxifen are shown as sticks and coloured yellow. **B**) X-ray crystal structure of the *rf*ERα LBD captured as a ‘head-to-head’ homodimer at 2.0 Å resolution (PDB: 9MDV). The structure reveals a novel orientation of H12 (shown in pink) which assumes a vertical orientation between H3 and H11, resulting in partial obscurement of the AF2. **C**) Molecular sidechain interactions between H3, H11 and H12 in apo (left) and active E2-bound states (right), highlighting formation of the hydrophobic cluster. Residue sidechains are shown as sticks and labelled. Highlighted in red are residues Y537 and D538 which are commonly mutated in ER+ breast cancer. **D**) Top-down perspectives of the ligand binding pocket (LBP) of apo (left) and E2-bound (right) structures showing side-by-side comparisons of pocket structure. Important residue sidechains are shown as sticks and labelled. E2 is shown as sticks and coloured yellow. **E**) Alignment of LBP residues of apo and active E2-bound structures. Apo residues are coloured blue, while active E2-bound residues are coloured green. E2 is shown as sticks and coloured yellow.

In the context of receptor activation, agonists of ERα, such as the primary sex hormone estradiol (E2), bind to the ligand binding pocket (LBP) of the ERα LBD and induce a structural rearrangement of H12, forming the activation function-2 (AF2) surface along with H3 – H5 (**Fig. 1A**). Once formed, the AF2 allows specific recruitment of coactivator proteins to the ERα LBD via a conserved *LxxLL* motif, leading to coactivation of transcription (*11*). Additionally, it has been identified that mutations within H12 of the ERα LBD such as Y537S and D538G, which are prevalent in ER+ breast cancer, can circumvent the requirement of agonist binding to form the AF2. This leads to constitutive activation of the receptor, driving tumour development and disease, and in some instances endocrine resistance (*2, 9, 12–15*).

Due to the role of ERα in breast cancer, extensive research has been performed to identify ligands which disrupt formation of the AF2 and repress ERα signalling. This has led to the discovery of frontline therapies termed selective estrogen receptor modulators/degraders (SERMs/SERDs), such as 4-hydroxytamoxifen, elacestrant and fulvestrant (*10, 16–20*). While SERMs/SERDs bind within the LBP they induce a different structural rearrangement, positioning H12 in a stable conformation that remodels the AF2 to antagonise transcriptional activity via several mechanisms including corepressor recruitment, immobilisation within the nucleus and degradation (*4, 21–25*). Although the structure of the ERα LBD in these active and repressed states has been thoroughly characterised (*4, 26–29*), there remains no structural information regarding the apo conformation of H12 in WT ERα that exists prior to ligand binding, or how H12 mutations such as Y537S and D538G escape this conformation to constitutively activate the receptor. Indeed, all present hypotheses for activation and inactivation of the receptor have been derived through retrospective analysis of ligand/peptide bound forms of the ERα LBD.

Here, we report a high-resolution crystal structure of an apo ERα LBD, capturing a third distinct H12 conformation that has not been previously observed. Combined with extensive biophysical and functional validation, this structure reveals that H12 functions as a ternary molecular switch that is modulated by ligand binding and destabilised by constitutively activating mutations, adopting three distinct states. Comparative structural and evolutionary analyses of ER and other nuclear receptors (NRs) challenges the notion of a universal activation mechanism. This study offers atomic-level insights into ligand-mediated modulation of a key transcription factor and the structural basis of cancer-driving mutations, with implications for therapeutic design.

## Results and discussion

### X-ray crystal structure of the apo *rf*ERα LBD reveals a unique H12 conformation

We initially attempted to structurally characterise the apo wildtype human ERα LBD, which has been historically intractable to crystallization. While a protocol to acquire pure, highly-concentrated apo protein (>40 mg/mL) was developed, we were unable to obtain crystals. Instead, we successfully solved the crystal structure of the wildtype apo ERα LBD from an Australian teleost fish, *Melanotaenia fluviatilis* (Murray River rainbowfish; *rf*ERα), previously studied by our laboratory (*30*), as a homodimer at 2.0 Å resolution (**Fig. 1B** and **Table S1**). Importantly, residues constituting the AF2 and H12 are highly conserved in *h*ERα and *rf*ERα orthologs. The crystal structure exhibits an overall global homology with the active E2-bound form (PDB: 9D8Q), with an RMSD of 0.775 Å between monomers. However, H12 adopts a conformation uniquely distinct from that of the active state (**Fig. 1C**). In the active state, H12 rests perpendicular to H3 and H4 forming the lower half of the AF2 interface. In contrast, in the apo state, H12 is orientated vertically and wedges between H3 and H11, enclosing the LBP and partially masking the AF2 with its C-terminal end (**Fig. 1C** and **fig. S1**). Analysis of the interaction between H12 and H3/H11 reveals that residues L536 and L540 of the H12 *LxxLL* motif interact with M343 and T347 (H3), W383 H5), and L525 (H11) to form a buried hydrophobic cluster. Externally, π-stacking between Y526 and Y537 and a salt bridge between K529 and D538G stabilise H12 in the vertical orientation. Together, these interactions shorten the H11-H12 loop by extending the α-helical structure of H12 and stabilise the apo conformation.

This distinct H12 conformation also results in significant changes within the LBP relative to the E2 bound receptor LBD (**Fig. 1D**). For apo *rf*ERα, the LBP of one monomer is occupied by three waters, and the other is occupied by a glycerol molecule, although no accessible solvent channels were observable. In the apo crystal structure, H11 and H12 undergo substantial restructuring compared to the active conformation. The vertical orientation of apo H12 pries H3 and H11 apart, with the latter rotating outward into the solvent. This movement causes H524 (which forms a hydrogen bond with the second hydroxyl group of E2) to flip out of the LBP, with its position instead occupied by L525 (**Fig. 1E**). Structural alignment of the two states reveals that E2 would clash with L525, L536, and L540 of the apo receptor. However, the residues and structure of H5, the β-sheet, H6 and H7 remain unchanged, except for E353 (H3) and R394 (H5), which form an ionic bond in the absence of ligand. The structural conservation of these residues between the two states suggests an important contribution to ligand binding by mediating coordination with the steroidal scaffold of the hormone. Taken together, these findings demonstrate that ligand binding physically modulates H12 conformation.

### H12 maintains a third discrete conformation in apo ERα LBD

To validate our apo crystal structure and suitability as a model for *h*ERα, we conducted biophysical experiments to compare the solution structure and dynamics of both LBDs. Native mass-spectrometry (native-MS) analysis of wildtype apo *rf*ERα and *h*ERα LBDs revealed that both receptors predominantly form homodimers consistent with the crystal structure, with a minor population of monomers also detected (**Fig. 2A**). Orthogonal validation using inline size-exclusion chromatography small-angle X-ray scattering (SEC-SAXS) revealed statistically similar scattering profiles between the two orthologs (CorMap *P* <0.01; Bonferroni corrected (*31*)), and SAXS molecular envelopes fitted to the crystal structure of the apo *rf*ERα LBD demonstrated conservation of homodimer architecture in solution, although the *h*ERα LBD was more compact (**Fig. 2B-C** and **fig. S2**; **Tables S2** and **S3**). No evident structural changes to the envelope encompassing H12 and surrounding area were observed between homologs.

**Figure 2.**
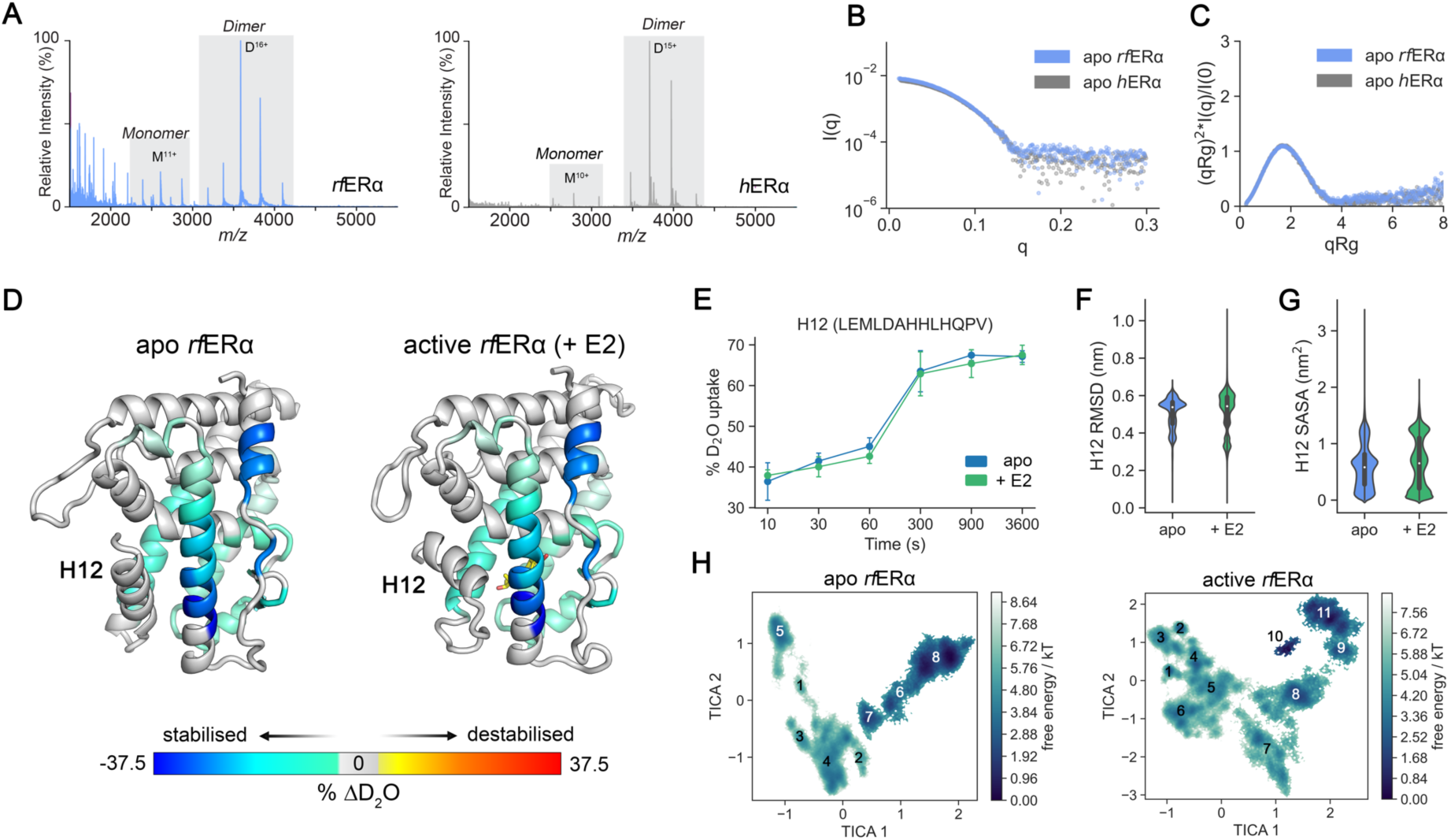
Comparison of the conformational landscapes of the hERα and rfERα LBDs. A) Native mass-spectrometry analysis of apo *rf*ERα and *h*ERα LBDs reveals the receptors predominantly form homodimers in the absence of estrogen (E2). **B**) Primary SAXS profiles of the *rf*ERα LBD (blue) and *h*ERα LBD (grey) represented as log I(q) plot versus q plot, and **C**) dimensionless Kratky plot demonstrating that protein samples are folded during SAXS data collection. **D**) Consolidated deuterium-uptake percentage differences (*Δ*D_2_O) between apo and active E2-bound *rf*ERα LBD obtained from hydrogen-deuterium exchange mass-spectrometry (HDX-MS) analysis mapped to structure. The data show that H1 and H3 undergo significant stabilisation upon E2 binding, alongside H5, the β-sheet, H7 and H11 to a lesser extent. Notably, deuterium uptake of H12 does not differ between states. **E**) Time-resolved D_2_O uptake of H12 (residues 541-553) for the apo *rf*ERα LBD and when bound to E2 at 10 s, 30 s, 60 s, 300 s, 900 s and 3600 s timepoints. Values represent the mean and standard deviation of at least two replicates. **F**) Violin plot comparing the distribution of average root-mean squared deviation (RMSD; in nm) of H12 backbone atoms between apo and active E2-bound states calculated from triplicate 5 µs all-atom molecular dynamics trajectories. **G**) As in **F**, but comparing average solvent accessible surface area (SASA; in nm^2^) of H12 residues. **H**) Re-weighted free energy landscapes (*G*/*kT*) of apo (left) and active (right) trajectories derived from Markov state modelling analysis of the ensembles. The free energy values (*G*/*kT*) are projected onto a time-structured independent component analysis (TICA) of backbone torsion angles. Approximate locations of metastable macrostates are labelled numerically.

To investigate changes in tertiary dynamics upon E2 binding, we performed HDX-MS analysis of the *rf*ERα LBD. Differential deuterium uptake revealed significant stabilisation (decrease in deuterium exchange) of H1, H3, the β-sheet, H6, H7, and the base of H11, upon E2 binding the apo receptor (**Fig. 2D**). In contrast, H12 showed minimal change (< 5%) in deuterium uptake/exchange (**Fig. 2E**), consistent with our structural data capturing H12 in two distinct, stable conformations representing the apo and E2-bound states of the *rf*ERα LBD (*30*). These results align with previous HDX-MS analyses of the *h*ERα LBD, which showed stabilisation of the same regions upon E2 binding, while the dynamics of H12 remained unchanged (*7–10*). All-atom molecular dynamics (MD) simulations of the wildtype *rf*ERα LBD demonstrated that the RMSD and solvent-accessible surface area (SASA) of H12 were comparable between the apo and active E2-bound state (**Fig. 2F** and **G**). Moreover, a comparison of average Cα atom root-mean squared fluctuation (RMSF) per residue agreed with the HDX-MS profile of the LBD, showing stabilisation of H1, H3, H5, β-sheet and H11 but no significant change in H12 dynamics (**fig. S3**). Comparison of H12 RMSD between apo *rf*ERα LBD trajectories and apo *h*ERα LBD trajectories (generated from a homology model) demonstrated that H12 remains relatively stable in the apo conformation with the *LxxLL* motif buried from solvent (**fig. S4**). Together, these data validate the apo crystal structure (show the agreement between the crystal structure and solution structure) and demonstrate that apo H12 exists in a third stable and discrete conformation, protecting buried residues from solvent exposure and regulating the accessibility of the AF2 surface to transcriptional coactivators.

### H3 allosterically modulates tertiary dynamics upon E2 binding

Agonist binding to the apo ER LBD induces widespread changes in tertiary dynamics, with H1, H3, the β-sheet, and H11 undergoing the most significant stabilisation determined by HDX-MS analysis (**Fig. 2D**). Yet, how these collective changes occur remains unclear. To address this, we performed additional E2-bound simulations of the *rf*ERα LBD, and then analysed both apo and E2-bound trajectories (30 µs total), with Markov State modelling (MSM) to resolve discrete ensembles and structural perturbations (**fig. S5** to **S7**) (*32, 33*). The analysis revealed that the apo and E2-bound LBDs traverse between distinct conformational landscapes, as described by backbone torsion angles (**Fig. 2H**). In the apo state, the receptor predominantly occupies one metastable state representing ∼80% of the stationary distribution, whereas the active E2-bound receptor mainly occupies three, similar low-energy metastable states representing ∼83% of the stationary distribution (**Table. S4** and **S5**). Structural analysis of metastable states show that the dynamic changes observed in the HDX-MS analysis primarily stem from conformational fluctuations of H3 prior to E2 binding. These analyses show that in the absence of ligand, H3 is dynamic and flexes in and out of the LBP functioning as a lever, distorting the conformation of H1 and the H1-H3 loop, H6, H7 and the β-sheet (**fig. S7**). Comparing between apo and E2-bound MSMs and the apo and E2-bound crystal structures shows that upon E2 binding, H12 is displaced, and expansion of the LBP by the hormone locks the H3 lever in the out conformation. E2 subsequently forms a dense network of interactions linking the β-sheet, H6 and H7, lowering their degrees of freedom. These findings indicate that E2 binding allosterically stabilises distal structural elements by modulating H3 conformation in addition to stabilising interactions with immediate residues, correlating with the HDX-MS analysis.

### Structural basis of constitutive activation

Our apo structural data provides an unprecedented opportunity to address longstanding questions regarding disease and ligand structure-activity relationships. In ER+ breast cancers, somatic mutations Y537S (and variants C/N/H) and D538G in H12 of the *h*ERα LBD constitutively activate the receptor, driving uncontrolled cellular growth and tumour development (*3, 9, 12, 34*). These mutations augment and confer receptor activation by introducing hydrogen bonding with D351 on H3 and improving H12 packing in the active conformation, respectively. The effects of these mutations on the apo receptor structure remain unclear. However, HDX-MS analysis of mutant *h*ERα LBDs show increased dynamics of the H11-H12 loop in the apo state compared to wildtype (specifically residues M528 to L540), suggesting that a destabilised H12 may more easily adopt a conformation amenable for coactivator binding in the absence of ligand (*9*). In the apo conformation, Y537 forms a π-stacking interaction with Y526, while D538 establishes a strong ionic bond with K529 (**Fig. 3A**). Structurally, the Y537S mutation disrupts the π-stacking by repelling Y529, while D538G severs the ionic bond with K529. Replicate MD simulations of apo mutant *rf*ERα and *h*ERα LBDs revealed notable destabilisation of residues spanning M528 to L540, consistent with the prior HDX-MS analysis (*9*). In the simulations, these mutations disrupt residue interactions leading to destabilisation of the H12-loop and a shift in the RMSD distribution compared to wildtype (**Fig. 3B**). We introduced the Y537S and D538G mutations into the *rf*ERα LBD to validate if constitutive activity is also conferred. Fluorescence polarisation coactivator recruitment assays showed a substantial increase in coactivator peptide binding relative to the wildtype apo receptor, and equivalently for the *h*ERα mutants (**Fig. 3C**), consistent with prior work (*9*). Together, these findings provide direct structural evidence demonstrating that oncogenic mutations Y537S and D538G disrupt critical interactions enabling escape of H12 from the apo conformation, allowing H12 to adopt the active conformation and constitutively activate transcription (**Fig. 3D**).

**Figure 3.**
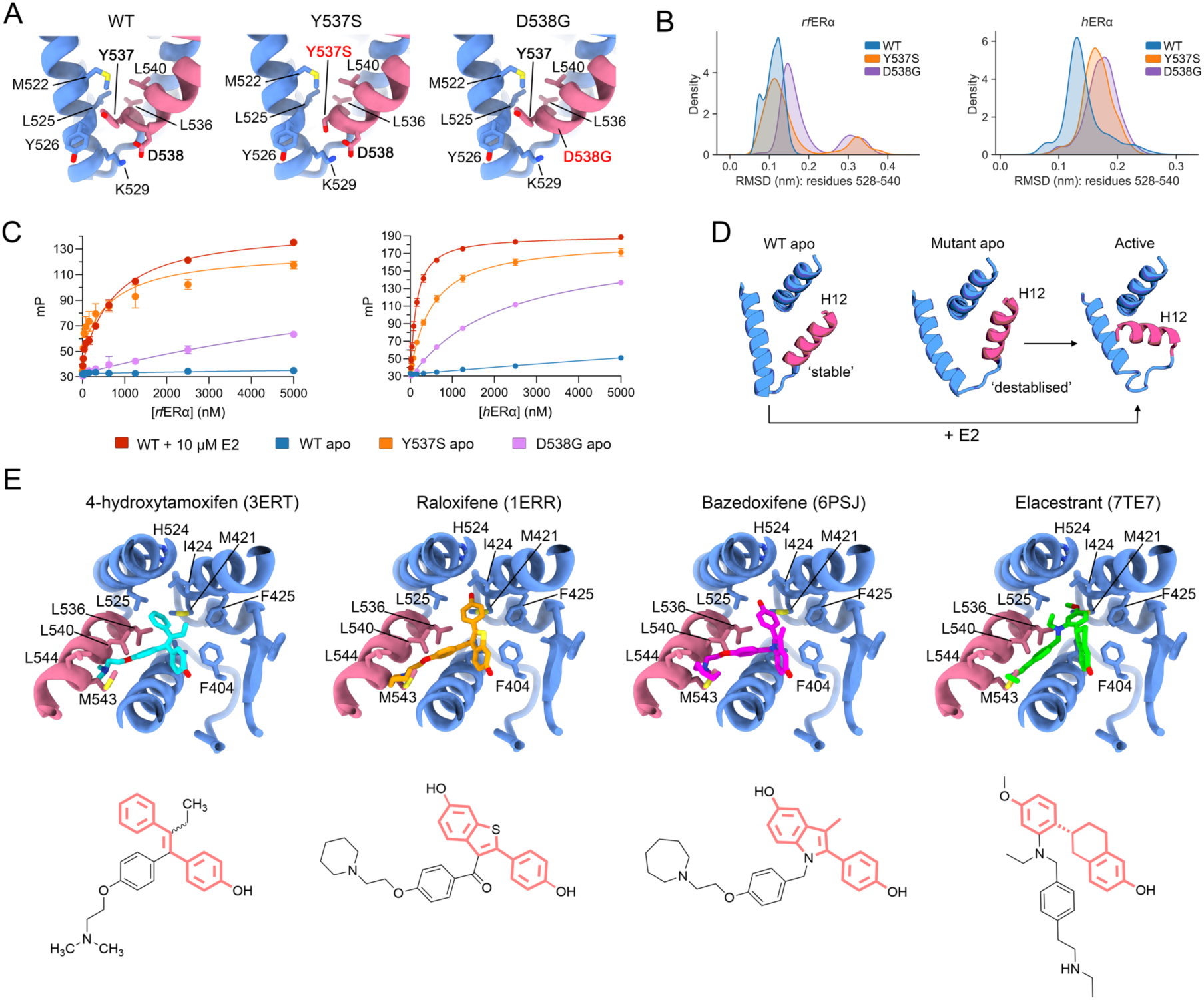
Structural basis of H12 escape and ligand-dependent antagonism. **A**) Oncogenic mutations Y537S and D538G disrupt key interaction that stabilise H12 in the apo conformation. Close-up perspective of the wildtype apo *rf*ERα LBD (left), and modelled Y537S (middle) and D538G (right) mutations. **B**) RMSD density distributions of the H11-H12 loop (residues 528-540) for wildtype, Y537S and D538G mutations, for apo *rf*ERα (left) and apo *h*ERα LBDs (right). **C**) I*n vitro* fluorescence polarization assays demonstrate coactivator recruitment for human an SRC-2 13-mer peptide probe (5-FAM–KHKILHRLLQDSS–COOH) for the *rf*ERα LBD (A) and *h*ERα LBD (right) with wildtype (± 10 µM E2), apo Y537S and apo D538G mutations. Error bars correspond to mean and standard deviation from three experiments. **D**) Cartoon schematic illustrating destabilisation of H12 in the apo conformation by Y537S and D538G mutations enabling escape to the active conformation in the absence of E2. **E**) (top) Crystallographic binding modes of the SERMs, 4-hydroxytamoxifen and raloxifene, and SERDs, bazedoxifene and elacestrant, from *h*ERα LBD crystal structures superimposed onto the apo *rf*ERα LBD crystal structure. PDB IDs are provided above each structure. Residue sidechains are shown as sticks and labelled. (below) Chemical structure of each ligand with core scaffold important for binding the apo receptor highlighted in pink.

### SERMs and SERDs displace H12 from the apo conformation

We then investigated how frontline SERMs and SERDs affect the apo ERα LBD conformation upon binding. Previous analyses of ligand-dependent inactivation have been hampered by the absence of a genuine apo structure. Understanding the key molecular interactions facilitating inactivation of the receptor upon initial ligand binding holds significant value for elucidating mechanisms of action and pharmacological development. Superimposing the apo *rf*ERα LBD with the inactive conformation *h*ERα LBD bound to the SERMs 4-hydroxytamoxifen and raloxifene, as well as the SERDs bazedoxifene and elacestrant, revealed that disruption of the apo conformation is primarily mediated by steric clashes between the ligand’s sidechains and H12 (**Fig. 3E**). Similar to E2 activation, each antagonist directly clashes with residues L525, L536, L540 and M543, displacing H12 into the solvent. However, the bulky chemical moieties of each ligand (e.g., the dimethylaminoethyl group of 4-hydroxytamoxifen) prevent H12 from adopting either the apo or active conformations and instead force H12 to bind within the partially-formed AF2 groove (*4, 26*). This mode of action is consistent with prior HDX-MS analyses of *h*ERα bound to SERMs and SERDs (*7–10*). We also observed that, akin to E2 binding the apo receptor, SERMs and SERDs make key interactions with residues within the LBP that are unchanged between states via a core scaffold (**Fig. 3E**). These findings suggest that small molecules which can facilitate displacement of the apo H12 conformation and share chemical similarity to the core scaffold may possess SERM- and SERD-like behaviour.

### A ternary molecular switch determines receptor function

Our findings reveal that H12 adopts a third discrete conformational state in the absence of ligand. We propose a model in which H12 functions as a ternary molecular switch that determines receptor activity (**Fig. 4A**). In this model, the receptor can exist in three distinct states where the output is determined by the chemical structure of the input ligand. In the first state (state 1), H12 adopts the apo conformation until a ligand binds to the receptor. If an agonist such as E2 binds, H12 transitions to the active conformation (state 2). Conversely, if a SERM or SERD such as 4-hydroxytamoxifen or elacestrant binds, H12 adopts the inactive conformation (state 3). The key event that triggers conformational switching, regardless of ligand, are steric clashes with L525 (H11), L536 and L540 (H12), which would destabilise and displace H12 into solvent. During transition to state 2, repositioning of H3 and H11 prevents H12 reverting to state 1, leading to the shielding of hydrophobic residues and the completion of the AF2 interface. However, in the presence of Y537S and D538G mutations, apo H12 is destabilised by loss of key contacts. This destabilisation lowers the threshold to state 2, allowing the receptor to adopt the active conformation. As a result, the ternary switch is no longer strictly controlled by ligand binding but is biased toward activation, enabling constitutive transcriptional activity. In contrast, during the transition to state 3, the SERM/SERD chemical structure prevents H12 reverting to state 1 or state 2, forcing it to occupy the AF2 groove via the helix’s *LxxLL* motif, ultimately repressing transcriptional activity.

**Figure 4.**
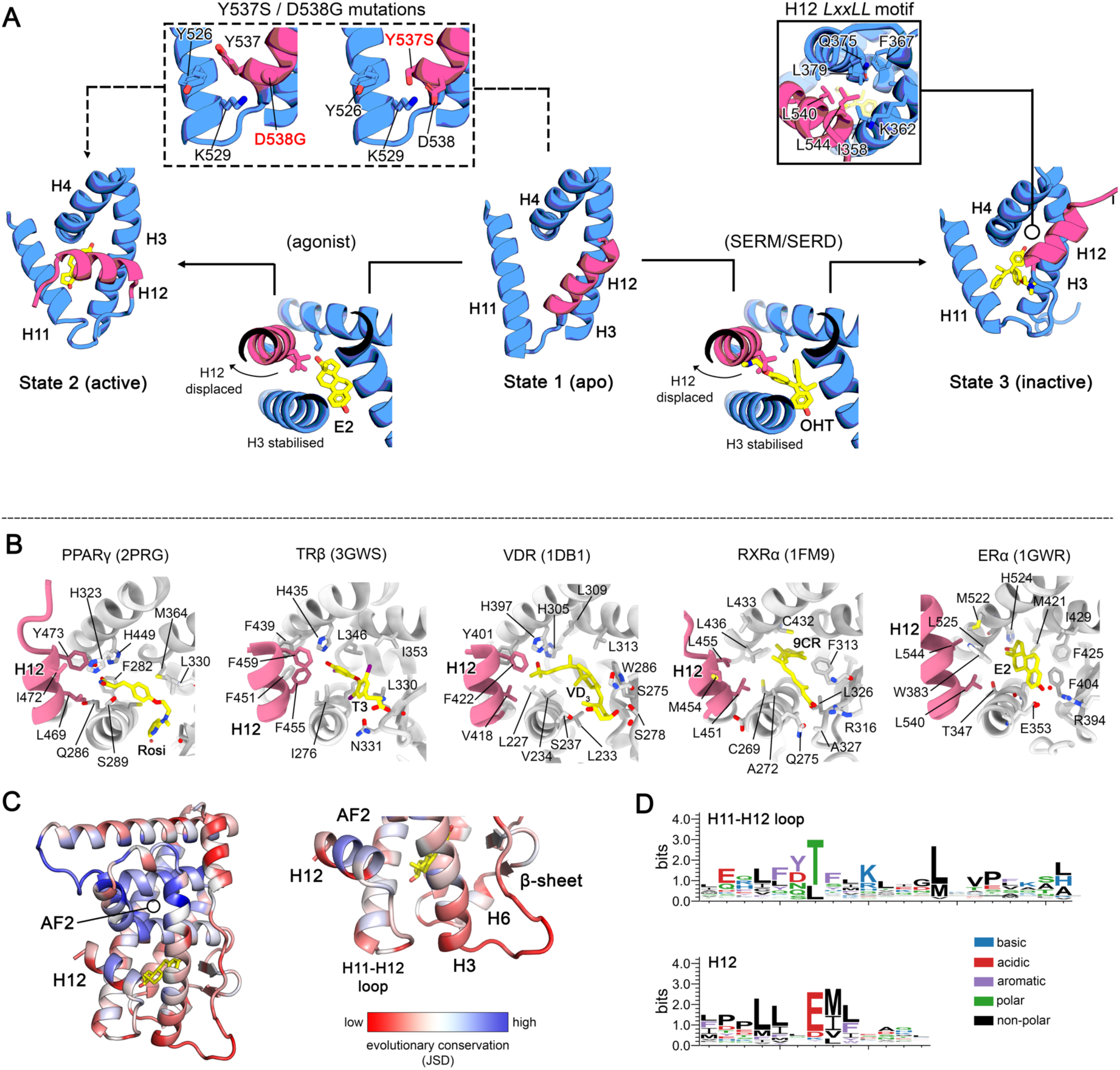
H12 functions as a ternary molecular switch to determine ERα LBD conformation. **A**) Cartoon diagram illustrating the ternary switch model. In the model, each state (1: apo; 2: active; 3: inactive) are shown as helices corresponding to H3, H4, H11 and H12 (in pink) for simplicity. Ligand binding to state 1 induces a conformational switch of H12 to either state 2 or state 3 but the outcome is dependent on the chemical structure of the ligand. However, in both outcomes the ligand stabilises H3 and clashes with the residues L525, L536 and L540 to displace H12 into solvent. In state 2, a requirement to shield hydrophobic residues from solvent, alongside repositioning of H3 and H11, stably anchors H12 in the active conformation thus forming the AF2. In state 3, the chemical structure of the SERM or SERD prevents H12 from reverting to the apo conformation or adopting the active conformation, instead causing the helix to stably bind in the partially formed AF2 binding cleft via its *LxxLL* motif. The oncogenic mutations Y537S and D538G disrupt key interactions enabling escape from the apo conformation and spontaneous transition to state 2 without the need for agonist binding. **B**) Top-down perspective showing ligand interactions within the LBP and H12 (in pink) for the LBDs of human PPARγ (PDB: 2PRG), TRβ (PDB: 3GWS), VDR (PDB: 1DB1), RXRα (PDB: 1FM9) and ERα (PDB: 1GWR). **C**) Evolutionary conservation of residue positions calculated as Jensen-Shannon divergence of 11,547 ligand-dependent NR LBD sequences projected onto the crystal structure of active E2-bound *h*ERα LBD. **D**) Sequence logo plots of the residue positions covering the base of H11 and the H11-H12 loop (top), and H12 (below) showing the sequence and chemical diversity of this region.

### Ligand-dependent activation mechanisms are diverse among NRs

The ternary switch model of ER activation is discordant to most other NRs which describes a dynamic ‘disorder-order’ transition of H12 upon ligand binding (*5, 6*). This mechanism is well-established for NRs like peroxisome-proliferator activated receptor γ (PPARγ), thyroid receptor (TR), and the vitamin D receptor (VDR) (*35–38*). Comparative structural analysis of ligand-bound NR LBDs demonstrates that H12 stabilisation mechanisms vary significantly. For example, in PPARγ, TRβ and VDR LBDs the ligands rosiglitazone, T3 and vitamin D directly contact H12 through hydrogen bonding networks and van der Waals interactions (**Fig. 4B**). Indeed, HDX-MS analysis and nuclear magnetic resonance (NMR) of these receptors show that H12 is significantly stabilised upon ligand binding, demonstrating a shift from either an unfolded or dynamic solvent-exposed ensemble (*36, 37, 39*). In contrast, ligands of retinoid-X receptor (RXR), ER and other steroid hormone receptors such as androgen receptor and glucocorticoid receptor, do not contact H12 due to structural differences in H3, H11 and the H11-H12 loop (**Fig. 4B**). In these receptors, H12 shields hydrophobic residues in the active conformation. Residues within this region are also important for conferring ligand specificity (*40*). For instance, between ER and AR the substitutions T347N (N705), H524F (F876) and L525T (T877) modify residues that are essential for E2 specificity and activation, instead contribute toward androgen hormone specificity in AR (*41*). Evolutionary analysis of more than 11,500 ligand-sensitive NR LBD sequences demonstrates that residues within this region are poorly conserved and vary in chemical properties compared to those which form the AF2 binding cleft (**Fig. 4C** and **D**). Consistent with this, the conservation of residues within the AF2 binding cleft would preserve the binding of coregulators (via the *LxxLL* motif) but in order to achieve stable interaction a functional H12 is necessary as mutation (or deletion) disrupts coactivation (*42–45*). Taken together, these findings indicate that ligand-activation mechanisms vary widely among nuclear receptors and likely coevolved alongside the emergence of ligand specificity to maintain regulatory functions.

## Conclusion

Here, we report the first crystal structure of an apo ER LBD and complemented by in-depth biophysical and computational analyses, reveal significant new insights into the mechanisms of ligand-dependent and -independent regulation. Our findings present a new model of receptor activation by hormones and inactivation by SERMs and SERDs used to treat ER+ breast cancer. The analysis revealed that both agonists and antagonist ligands disrupt apo H12 conformation through clashes with L525, L536 and L540, and coordinate with residues in the LBP via a similar core chemical scaffold. We also demonstrate that oncogenic mutations Y537S and D538G disrupt crucial contacts that stabilise the apo conformation of H12 providing a structural basis for their constitutive activating function in ER+ breast cancer. In summary, this work uncovers new insight into the regulation of a key transcription factor and serves as a foundation to inform new therapeutic design.

## Materials and Methods

### Protein expression and purification

#### Construct design

Codon-optimised inserts for the wildtype ligand binding domains (LBDs) of *Melanotaenia fluviatilis* ERα (*rf*ERα) and human ERα (*h*ERα), and also for Y537S and D538G variants, were cloned into a pET-11a expression vector with ampicillin resistance and synthesised by GenScript Biotech (Piscataway, NJ, USA). The *rf*ERα inserts contained an N-terminal non-removable hexahistidine tag, while the *h*ERα inserts contained a removable N-terminal hexahistidine + SUMO solubility tag. A sequence alignment comparing each insert is provided in Supplementary figure 2.

#### Expression of apo rfERα and hERα LBDs

Expression plasmids were transformed into either *Escherichia coli* strains BL21(DE3) (apo) or LEMO21(DE3) (Y537S and D538G). Initially, 3 x 1 L conical flasks containing Luria broth (LB) supplemented with 100 µg/mL ampicillin were inoculated with 20 mL (1:50 dilution) of an overnight seed culture transformed with the expression plasmid of interest, and grown at 37 °C with shaking (200 rpm) for aeration. Upon the culture reaching an optical density of A_600 nm_ value of 0.8 – 1.0, expression of recombinant protein was induced by addition of IPTG to a final concentration of 500 µM at 16 °C for 20 hours at 200 rpm. The expression protocol for *rf*ERα LBDs and *h*ERα LBDs were identical, except for the host strain as described above.

#### Purification of apo rfERα LBDs

All purification steps described herein were performed at 4 °C or on ice. Following overnight expression, cells were harvested by centrifugation at 5000*g* for 10 minutes and each pellet was resuspended in 25 mL of ice-cold Buffer A (20 mM Tris-HCl pH 8.0, 500 mM NaCl, 10 mM imidazole, 2 mM β-mercaptoethanol, and 0.1% Tween 20) and lysed by mechanical disruption. The lysates were clarified by centrifugation at 40,000*g* for 30 minutes, and then loaded onto a 5 mL HisTrap HP column (GE Life Sciences) connected to an NGC Medium-Pressure Liquid Chromatrography System (Bio-Rad) at a flow rate of 4 mL/min, equilibrated with Buffer B (20 mM Tris-HCl pH 8.0, 500 mM NaCl, 10 mM imidazole, 2 mM β-mercaptoethanol). The column was washed with 10 column volumes of Buffer B and then with 5 column volumes of 20% Buffer C (100% = 20 mM Tris-HCl pH 8.0, 500 mM NaCl, 250 mM imidazole, 2 mM β-mercaptoethanol) to remove weakly bound impurities. Bound *rf*ERα LBD was eluted in two steps of 5 column volumes of 65% Buffer C and then 5 column volumes of 100% Buffer C. The elution fractions containing *rf*ERα LBD were identified by SDS-PAGE analysis diluted ∼8-fold into no-salt Buffer D (20 mM Tris-HCl pH 8.0, 10% glycerol, 5 mM DTT). The diluted sample (∼200 mL in total), was immediately loaded onto a 5 mL HiTrap Q HP ion exchange column (GE Life Sciences) equilibrated in Buffer E (20 mM Tris-HCl pH 8.0, 100 mM NaCl, 10% glycerol, 5 mM DTT) at a flow rate of 4 ml/min. The column was then washed with 5 column volumes of Buffer E and the bound protein was eluted over a 20-column volume linear gradient of 0 – 100% Buffer F (20 mM Tris-HCl pH 8.0, 500 mM NaCl, 10% glycerol, 5 mM DTT). The bound *rf*ERα LBD typically eluted at ∼ 200 mM NaCl concentration. The elution fractions containing highly pure *rf*ERα LBD were pooled and concentrated to ≥ 10 mg /mL using an Amicon Ultra centrifugal filter (10,000 MW cut-off, Sigma Aldrich) and flash-cooled in liquid nitrogen prior to storage at -80 °C. Protein concentrations were determined by measuring the absorbance at 280 nm and using the molar extinction coefficient and molecular weight calculated by ProtParam (https://web.expasy.org/protparam/).

#### Purification of apo hERα LBDs

The purification of *h*ERα LBDs by nickel affinity and anion exchange chromatography were performed as described above for *rf*ERα LBDs, with the exception that Buffer A was replaced with Buffer B, and Buffers D and E contained 2 mM β-mercaptoethanol in place of 5 mM DTT. Following the anion exchange chromatography step, elution fractions containing SUMO-*h*ERα LBD were pooled and incubated with 0.5 mg of recombinant SUMO protease overnight at 4 °C. The digested sample was then loaded onto a 5 mL HisTrap HP column (GE Life Sciences) equilibrated in Buffer E (20 mM Tris-HCl pH 8.0, 500 mM NaCl, 10 mM imidazole, 2 mM β-mercaptoethanol). The digested *h*ERα LBD was eluted by washing the column with 5 column volumes of 10% Buffer C, with the His-SUMO and SUMO protease staying bound to the column. Elution fractions containing highly pure *h*ERα LBD were pooled and buffer exchanged 200-fold into Buffer E using an Amicon Ultra centrifugal filter (10,000 MW cut-off, Sigma Aldrich) and flash-cooled in liquid nitrogen prior to storage at -80 °C. The concentrations of *h*ERα LBDs were determined using the same method as *rf*ERα LBDs.

### Fluorescence polarisation coactivator recruitment assays

The human SRC2-2 coactivator peptide probe (5-FAM–KHKILHRLLQDSS–COOH; 98% purity) was purchased from GenScript Biotech (Piscataway, NJ, USA) (*46*). For fluorescence polarization assays a 200 µM estradiol stock was prepared in 100% DMSO. The coactivator probe was prepared as a 10 mM stock in MQ H_2_O and adjusted to ∼pH 7 with dropwise addition of 2M NaOH, with working stocks subsequently prepared at a final concentration of 100 µM in 10 mM Tris-HCl pH 7.5 and stored at -20°C.

All fluorescence polarisation experiments were performed in triplicate as previously described with slight modification (*47*). Under apo conditions, samples (at 160 µL final volume) contained FP buffer (20 mM Tris-HCl pH 8.0, 100 mM NaCl, 0.05% Tween 20, 1 mM TCEP, 5% DMSO, 10% glycerol and 50 nM SRC2-2 probe), with the protein concentrations of each ER LBD variant varied. For WT *h*ERα and *rf*ERα LBDs, the experiment was also performed in the presence of 10 µM E2. Once prepared, the samples were incubated at 22 °C for 30 minutes, and then 150 µL of each sample transferred to a single well of a non-binding flat-bottom black 96 well plate (Greiner Ref. 655900). Fluorescence polarization was measured using a Pherastar FS microplate reader equipped with the FP 485 520 520 module, using default settings. mP values were calculated using Equation 1 following subtraction of raw parallel and perpendicular emission values for a corresponding sample with the coactivator probe omitted.

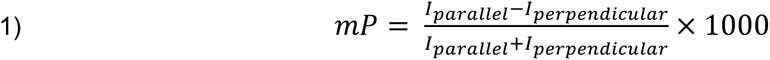

To investigate the association between the coactivator class probe with ER LBDs in the presence and absence of E2, the samples comprised FP Buffer and 8 different concentrations of ER LBD ranging between 0.3 and 5× the estimated dissociation constant (Kd). This was estimated by non-linear regression in GraphPad Prism using Equation 2, where *mP_free_* and *mP_bound_* are the fluorescence polarization for free probe and saturated receptor respectively, [*L*] is the total concentration of coactivator class probe, [*R*] is the total concentration of ER LBD homolog and *K_d_* is the dissociation constant.

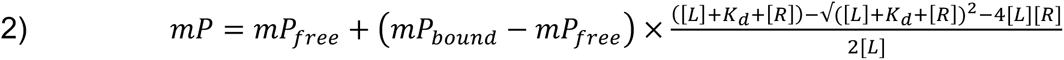

### X-ray crystallography

Crystallisation experiments were performed using the sitting drop vapour diffusion method in 96-well Intelliplates (ArtRobbins) at 16 °C. Conditions promoting crystal formation were probed using the NR-LBD, PEG/Ion HT, Index HT, and Crystal Screen HT sparse matrix screens (Hampton Research). Reservoir volumes were 80 µL and crystals were formed in drops containing 1 µL 16 mg/mL apo *rf*ERα LBD to 1 µL of reservoir solution. Diffracting crystals were obtained in the NR-LBD sparse matrix screen condition E5 (0.1 M NaCl, 0.1 M HEPES pH 7.0, and 225 PEG 2000 MME), appearing as large rods (∼500x50x50 µm) after 1 week.

### Data processing and refinement

Prior to data collection, crystals were cryoprotected by immersion in NVH oil and flash cooled in liquid nitrogen. Crystals were diffracted at 100 K at the MX1 beamline of the Australian Synchrotron (*48*). Images were indexed and integrated in XDS, with scaling and merging completed in Aimless (*49, 50*). The phase problem was solved by molecular replacement in Phaser using Chain A of a previously determined structure of the *rf*ERα LBD complexed with E2 and a human SRC2-2 coactivator-derived peptide (PDB: 9D8Q) as the search model, with ligands and solvent removed (*51*). The resulting model underwent several rounds of refinement in phenix.refine and rebuilding in Coot, until the refinements converged (*52, 53*). The final model displayed good overall geometry and contained no Ramachandran outliers. Data collection and refinement statistics are detailed in **Table S1**.

### Native mass-spectrometry

Protein samples containing wildtype *h*ERα or *rf*ERα LBDs were diluted into 15 mL of ammonium bicarbonate pH 7.0 and buffer exchanged five times using an Amicon Ultra centrifugal filter (10,000 MW cut-off, Sigma Aldrich) to a final concentration of 10 µM. Native mass-spectrometry was performed on a Bruker Impact II HDMS Q-ToF mass spectrometer (Bruker Daltonics) with a nanoelectrospray ionisation source with the following instrument parameters: *m/z* range, 500-10,000; polarity, positive; capillary voltage, 2 kV; end plate offset, 500 V; and source temperature, 60 °C. Protein samples were electrosprayed from platinum-coated borosilicate glass capillaries (Harvard Apparatus, USA) prepared in-house. All spectra were analysed using the Compass Data Analysis software (v4.2 Bruker Daltonics). Protein masses were manually deconvolved using *m/z* and charge values.

### Hydrogen-deuterium exchange detected by mass-spectrometry

#### Peptide identification

Differential HDX-MS experiments were conducted as previously described with several modifications (*54*). Peptides were identified using tandem MS (MS/MS) with an Orbitrap mass spectrometer (Q Exactive, ThermoFisher). Product ion spectra were acquired in data-dependent mode with the top five most abundant ions selected for the product ion analysis per scan event. The MS/MS data files were submitted to Mascot (Matrix Science) for peptide identification. Peptides included in the HDX analysis peptide set had a MASCOT score greater than 20 and the MS/MS spectra were verified by manual inspection. The MASCOT search was repeated against a decoy (reverse) sequence and ambiguous identifications were ruled out and not included in the HDX peptide set.

#### HDX-MS analysis

5 μl of 10 µM of apo *rf*ERα LBD, or complexed with 5:1 molar ratio E2, was diluted into 20 μl D_2_O buffer (20 mM Tris, pH 7.8; 150 mM NaCl; and 2mM DTT) and incubated for various time points (0, 10, 60, 300, 900 and 3600s) at 4°C. The deuterium exchange was then slowed by mixing with 25 μl of cold (4°C) 3 M urea, 50mM TCEP, and 1% trifluoroacetic acid. Quenched samples were immediately injected into the HDX platform. Upon injection, samples were passed through an immobilized pepsin column (1mm × 2cm) at 50 μl min^−1^ and the digested peptides were captured on a 1mm × 1cm C_8_ trap column (Agilent) and desalted. Peptides were separated across a 1mm × 5cm C_18_ column (1.9 μl Hypersil Gold, ThermoFisher) with a linear gradient of 4% - 40% CH_3_CN and 0.3% formic acid, over 5 min. Sample handling, protein digestion and peptide separation were conducted at 4°C. Mass spectrometric data were acquired using an Orbitrap mass spectrometer (Exactive, ThermoFisher). HDX analyses were performed in triplicate, with single preparations of each complex form. The intensity weighted mean *m/z* centroid value of each peptide envelope was calculated and subsequently converted into a percentage of deuterium incorporation (%D). This is accomplished determining the observed averages of the undeuterated and fully deuterated spectra and using the conventional formula described elsewhere (*55*). Statistical significance for the differential HDX data is determined by an unpaired t-test for each time point, a procedure that is integrated into the HDX Workbench software (*56*). Corrections for back-exchange were made on the basis of an estimated 70% deuterium recovery, and accounting for the known 80% deuterium content of the deuterium exchange buffer. Each %D is averaged across all time points to determine an overall %D for each peptide.

#### Data rendering

The HDX data from all overlapping peptides were consolidated to individual amino acid values using a residue averaging approach. Briefly, for each residue, the deuterium incorporation values and peptide lengths from all overlapping peptides were assembled. A weighting function was applied in which shorter peptides were weighted more heavily and longer peptides were weighted less. Each of the weighted deuterium incorporation values were then averaged to produce a single value for each amino acid. The initial two residues of each peptide, as well as prolines, were omitted from the calculations. This approach is similar to that previously described (*57*).

### Size-exclusion chromatography coupled small-angle X-ray scattering

Small-angle X-ray scattering experiments were performed on the BioSAXS beamline on the Australian Synchrotron. Concentrated protein samples were diluted into buffer containing 20 mM Tris-HCl pH 7.5, 150 mM NaCl, 5% glycerol and 2 mM TCEP and buffer exchanged five times using an Amicon Ultra centrifugal filter (10,000 MW cut-off, Sigma Aldrich) to a final concentration of 5 mg/mL. The samples were loaded onto an equilibrated S200 5/150 GL column (Cytiva) using the Coflow autoloader (*58*) at a flow rate of 0.3 mL/min through a 3 mM UV flow cell (Knauer, Berlin, Germany). UV absorbances at 260 and 280 nm were measured immediately before presentation to the X-ray beam by an STS microspectrometer (Ocean Optics, Orlando, FL, USA). X-ray scattering was measured continuously during SEC at 293K with a 1 s exposure time at 12.4 keV (λ = 1.00 Å) using a Pilatus3X 2M detector positioned 2,150 mm (1,500 mm + 650 mm offset distance) from the sample capillary. This afforded a q-range of 0.0098 to 0.72027 Å^-1^ following data reduction. Data was reduced by Fast Azimuthal Integration using Python and PyFAI with customised algorithms written for the BioSAXS beamline. Data were placed on an absolute scale using water in the measurement capillary as a standard and the nominal diameter of the capillary at the measurement position was 1.0 mm. The subtracted profiles were analysed using BioXTAS RAW (v2.3), and GNOM implemented in ATSAS (v3.2.1) (*59, 60*). SAXS envelopes were calculated using the DAMMIF/N algorithm implemented in ATSAS and fit to the high-resolution crystal structure of the apo *rf*ERα LBD.

### Molecular dynamics (MD) simulations

All-atom MD simulations for wildtype *rf*ERα (+/- E2) and apo *h*ERα LBDs, along with Y537S and D538G variants, were performed using the same method outlined here, except for the duration of production simulations. MD simulations were performed using GROMACS with the Charmm36 forcefield (*61–64*). E2 topology was generated using SwissParam (*65*). Protein coordinates were placed in the centre of a dodecahedral box, situated at least 10 Å from the from the periodic edge boundary. The systems were solvated with the TIP3P water model and to neutralise net charge, Na^+^ and Cl^-^ ions were added where required. Following system preparation, energy minimisation was performed using the steepest descent method for a maximum of 50,000 steps or until *F_max_* reached < 1000 kJ/mol/nm. Energy minimised systems were equilibrated using sequential 1 ns restrained simulations in the NVT and NPT ensembles at 310 K using Lincs constraint algorithm (hydrogen bonds), Particle Mesh Ewald electrostatics, Berendsen thermostat coupling, and for the NPT ensemble only, Parrinello-Rahman pressure coupling. Restraints were then released and production simulations performed in the NPT ensemble at 310 K using a 2-femtosecond timestep, with frames stored every 100 ps. Production simulations were performed in triplicate for at least 1 µs duration. Analyses were performed in Python using the MDTraj, NumPy, Matplotlib and Seaborn libraries (*66–69*).

### Markov modelling

To structurally analyse the conformational ensembles, Markov state modelling was employed; a powerful approach for studying kinetic data generated by MD trajectories, implemented with PyEMMA (*32, 70*). Feature space was defined by the backbone torsion angles of the receptor, which were then discretised into two dimensions using time-structured independent component analysis (TICA). Microstate clustering was performed on the first two TICA components using the K-means algorithm, different cluster sizes were explored to determine the optimal number microstate centres based on convergence of the implied timescale (ITS) plots. Ultimately, *k*=250 was chosen for the apo state and *k*=400 was chosen for the active E2-bound state. The ITS plots resolved 7 slow processes at a lag time of 1 ns for the apo state and 10 slow processes at a lag time of 5 ns for the active state. Bayesian Markov State Models (MSMs) were built and statistically validated using the Chapman-Kolmogorov test. To coarse grain the models into macrostates, Perron-cluster cluster analysis (PCCA) was used and transition-path theory (TPT) was implemented to estimate mean-first passage times (MFPTs). Microstate assignments were extracted from the models by sample distribution and analysed with PyMOL (Schrödinger, LLC).

### Evolutionary sequence analysis

Primary amino acid sequences for ligand-activated NR LBDs were curated from the UniRef-100 and UniRef-90 databases with *jackhmmer* using the human sequence as a query (*71*). Assembly of the multiple sequence alignment was performed in a three-step process. First, hits from *jackhmmer* queries of each receptor were aligned in MAFFT and edited to human residue positions in order to remove lineage-specific indels. Sequences with more than 30% gaps were removed. Following this, human NR LBD sequences were iteratively aligned with MAFFT to create a reference sequence profile which was used to merge the individual alignments using *hmmalign* software. To calculate the evolutionary conservation, the Jensen-Shannon divergence (JSD) of each residue position was calculated with SciPy, using the background (*q*) and position-specific (*p*) amino acid probability distributions (*72*). Sequence logos were created using WebLogo3 (https://weblogo.threeplusone.com/) (*73*).

## Acknowledgements

This research was undertaken in part using the MX1 beamline at the Australian Synchrotron, part of ANSTO, and made use of the Australian Cancer Research Foundation (ACRF) detector. Computational resources were provided by the University of Adelaide Phoenix High-Performance Computing (HPC) facility. D.P.M. was supported by an Australian Government Research Training Program (RTP) scholarship. B.J. is supported by The Hospital Research Foundation Group Research Fellowship. This work is supported by an Australian Research Council Discovery Project (DP230100609).

## Author contributions

Conceptualization: D.P.M., J.L.P., J.B.B. Funding acquisition: J.B.B. Methodology: D.P.M., J.L.P. S.N., B.J., A.M. Investigation: D.P.M., J.L.P. S.N., B.J. Data analysis: D.P.M., J.L.P., S.N., B.J. Software: D.P.M. Visualization: D.P.M. Writing – original draft: - D.P.M. and J.L.P. Writing – review and editing: - all authors.

## Data and material availability

Atomic coordinates for the X-ray crystal structure of the apo *rf*ERα LBD are deposited in the Protein Data Bank under the accession code 9MDV. SEC-SAXS data generated in this study are deposited in the Small Angle Scattering Biological Data Bank (SASBDB) under accession IDs: SAS6626 (apo *h*ERα LBD) and SAS6628 (apo *rf*ERα LBD).

