## Supplementary information for "A ternary switch determines ERα LBD conformation"

**The file includes:**

Materials and Methods

Figs. S1 to S7

Tables S1 to S7

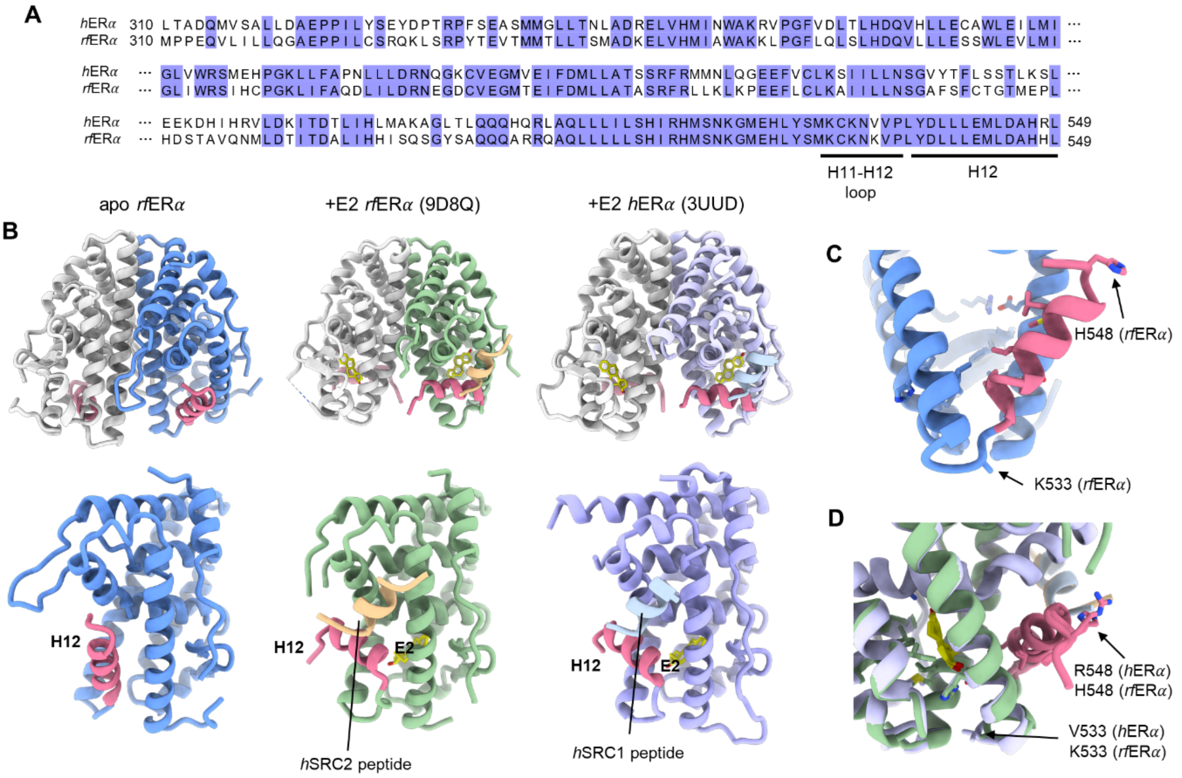


**Fig. S1.** X-ray crystal structure of the apo *rf*ERα ligand binding domain (LBD). **A**) Sequence alignment comparing the LBD of *h*ERα (top) and *rf*ERα (bottom) LBDs. Highlighted residues are those which are conserved. The H11-H12 loop and H12 are underlined. **B**) X-ray crystal structures of the apo *rf*ERα LBD (this study), active E2-bound *rf*ERα LBD (PDB: 9D8Q) and active E2-bound *h*ERα LBD (PDB: 3UUD) shown as the homodimer (top) and monomer (below). H12 is highlighted in pink and coactivator peptides are shown. E2 is shown as sticks and coloured yellow. **C** and **D**) The two residues which differ with the H11-H12 loop and H12 of both receptors, V/K533 and R/H548, shown in the apo *rf*ERα LBD structure and active E2-bound structures of *rf*ERα and *h*ERα LBDs (green and blue, respectively), do not form important interactions with other residues, if any. Note that in both apo and active E2-bound *rf*ERα LBD structures K533 is not modelled in the electron density.

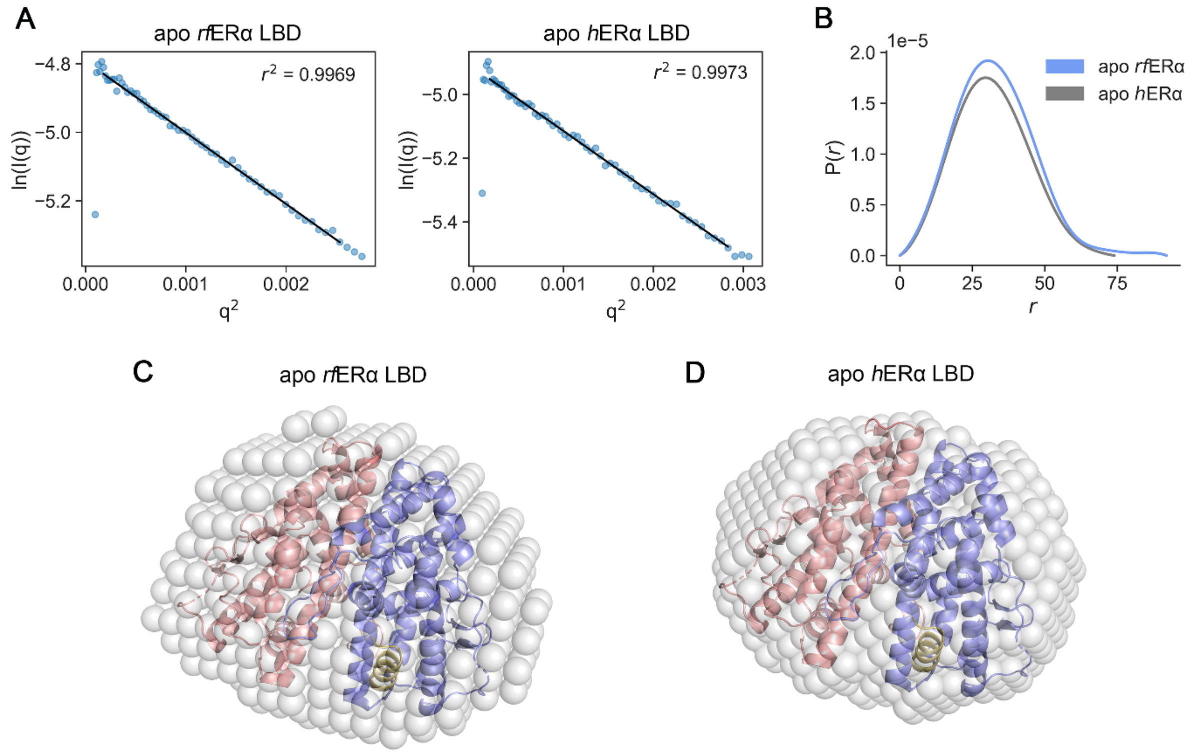


**Fig. S2**. **A**) Guinier plots of the SEC-SAXS data representing the log linear scattering intensity given as ln(I(q)) versus the scattering vector, q, for both experiments. The plots show the high quality of the scattering data, with minimal aggregation and interparticle repulsion. Radius-of-gyration (Rg) of the apo *rf*ERα LBD was calculated to be 25.0 ± 0.0797 Å; and Rg of the apo *h*ERα LBD was calculated as 24.42 ± 0.0783 Å. **B**) Pair distance distribution function plot of the scattering data. **C**) Refined DAMMIF/N model of the apo *rf*ERα LBD SAXS data fit to the corresponding high-resolution crystal structure (PDB: 9MDV) ($\chi^2$ = 1.215). **D**) Refined average DAMMIF/N models of the apo *h*ERα LBD SAXS data fit to the high-resolution crystal structure of the apo *rf*ERα LBD (PDB: 9MDV) ($\chi^2$ = 1.222). In both **C** and **D**, the SAXS envelope is shown as transparent grey spheres, and the structural model is represented as a cartoon. Chain A is coloured blue and chain B is coloured pink; H12 is highlighted in yellow. While the models clearly show conservation of homodimer architecture and no evident differences around the H12 region, the apo *h*ERα envelope is slightly more compact.

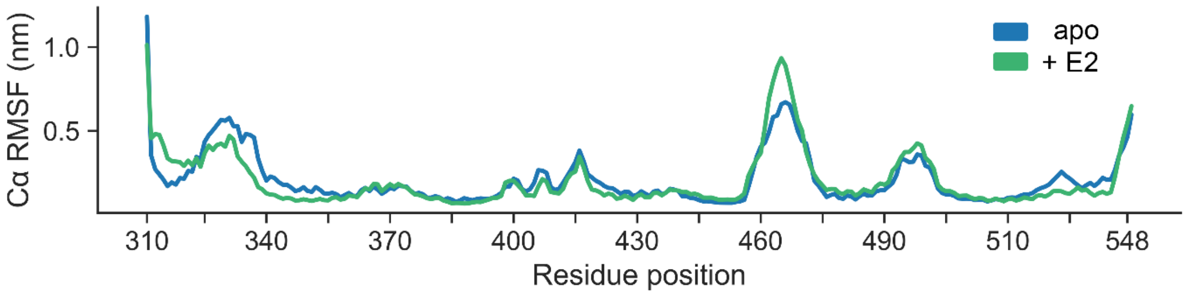


**Fig. S3**. Average root mean-squared fluctuation (RMSF) of Cα atoms from three replicate 5 μs duration all-atom molecular dynamics simulations for apo (blue) and E2-bound *rf*ERα LBD.

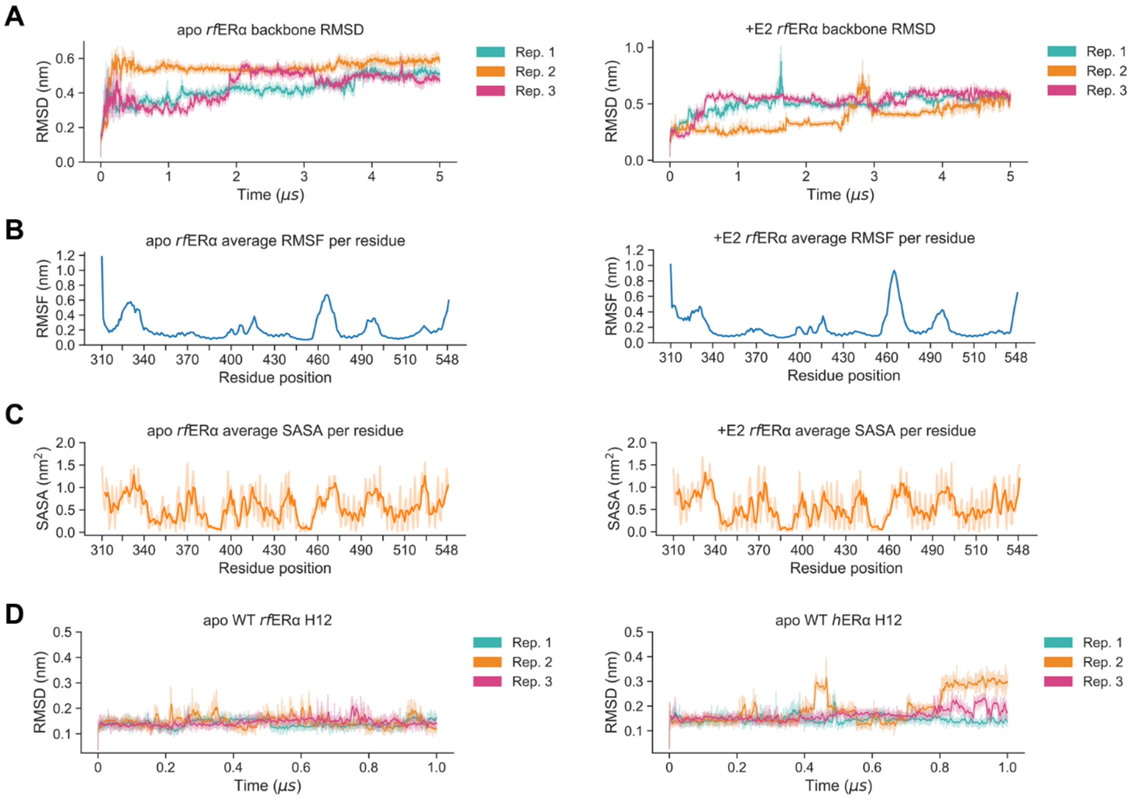


**Fig. S4.** All-atom molecular dynamics (MD) analysis of the apo and active E2-bound *rf*ER$\alpha$ LBD. **A**) Heavy backbone atom root-mean squared deviation (RMSD) compared to starting coordinates for each replicate 5 $\mu$s trajectory. The solid line represents a 5 ns rolling average, superimposed atop the raw RMSD values. **B**) Average C$\alpha$ root-mean squared fluctuation (RMSF; in nm) per residue from the three replicate 5 $\mu$s trajectories. **C**) Average solvent-accessible surface area (SASA; in $nm^2$) per residue from the three replicate 5 $\mu$s trajectories. The solid line shows a three-residue rolling average. **D**) Heavy backbone atom RMSD of H12 residues (537-546) over the course of three 1 $\mu$s trajectories for apo *rf*ERα (left) and *h*ERα (right).

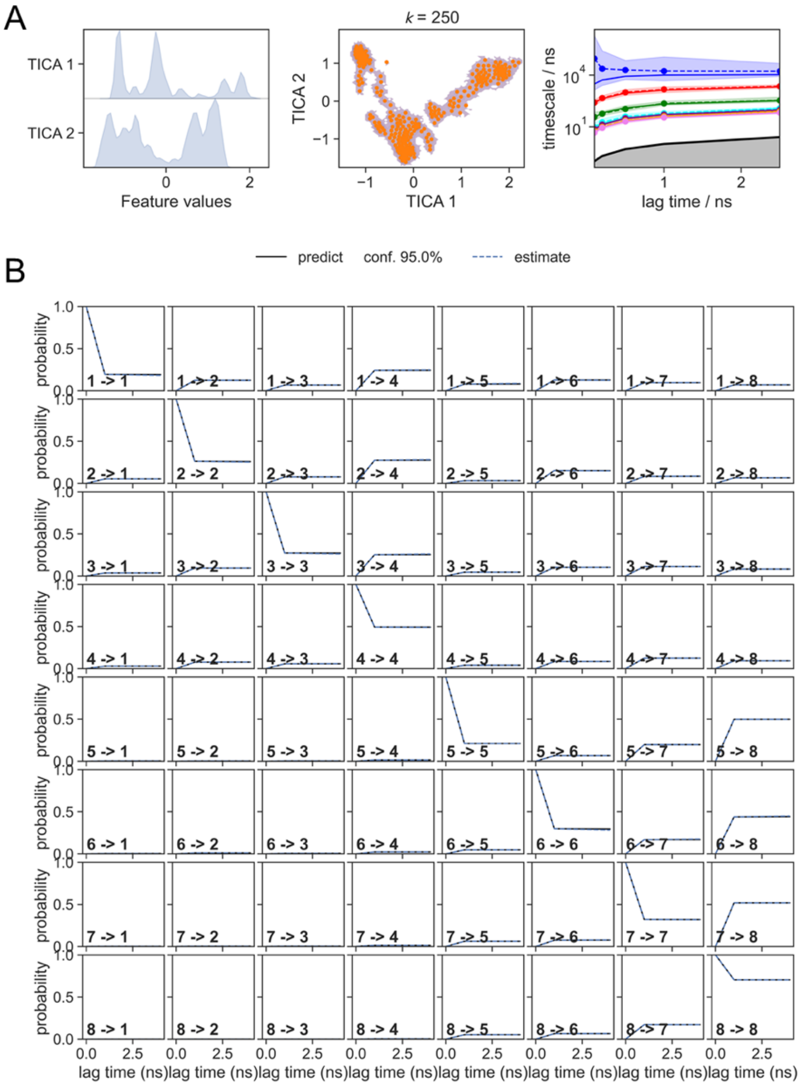


**Fig. S5**. Implied timescale plot and statistical validation of the apo *rf*ER*α* Markov model using the Chapman-Kolmogorov test. **A**) Left: Distribution of backbone torsion angles ("Feature values") in the space of the first two components from the time-structured independent component analysis (TICA). Middle: K-means cluster centroids (microstates) mapped to coordinates of the first two TICA components. Right: Implied timescales (ITS) plot using *k* = 250 microstates shows convergence of 7 slow-processes (8 metastable states) at a lag time of 1 ns. **B**) Statistical validation of the Bayesian Markov model estimated using 8 metastable states and a lag time of 1 ns.

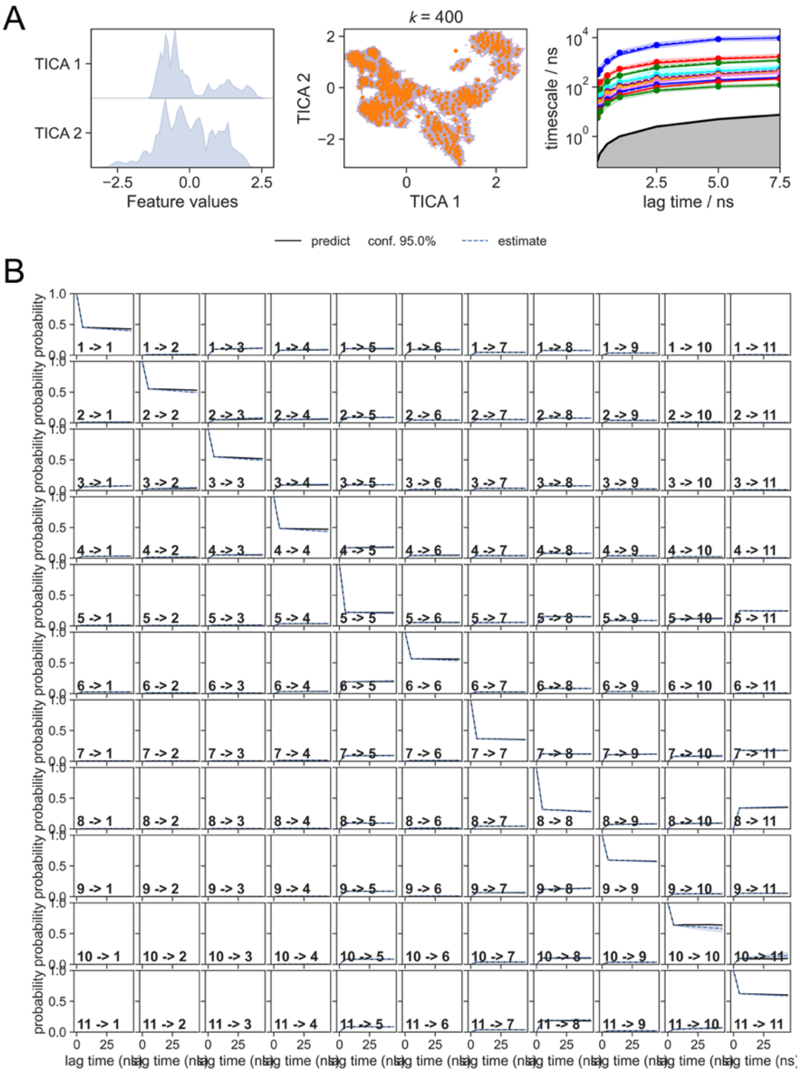


**Fig. S6**. Implied timescale plot and statistical validation of the active E2-bound *rf*ER$\alpha$ Markov model using the Chapman-Kolmogorov test. **A**) Left: Distribution of backbone torsion angles ("Feature values") in the space of the first two components from the time-structured independent component analysis (TICA). Middle: K-means cluster centroids (microstates) mapped to coordinates of the first two TICA components. Right: Implied timescales (ITS) plot using $k$ = 400 microstates shows convergence of 7 slow-processes (8 metastable states) at a lag time of 1 ns. **B**) Statistical validation of the Bayesian Markov model estimated using 8 metastable states and a lag time of 1 ns.

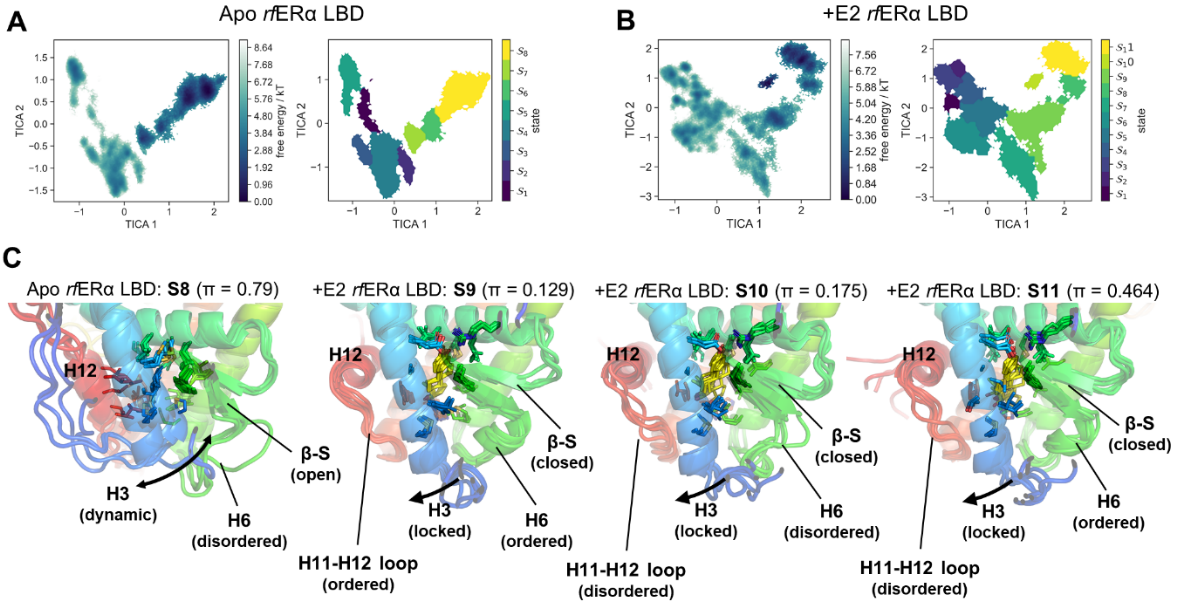


**Fig. S7.** Re-weighted free energy surfaces (left) and coarse-state map (right) plotted onto TICA reduction coordinates of the apo *rf*ERα LBD (**A**) and +E2 *rf*ERα LBD (**B**). **C**) Top five microstates sampled by distribution from the apo *rf*ERα LBD MSM (state 8), and +E2 *rf*ERα LBD MSM (states 9 to 11). The stationary distribution of each state (π) is included in the header. In the apo state, H3 transitions between an 'in' and 'out' conformation within the LBP. This destabilises the end of H5, the β-sheet (in an open conformation), H6 and H7. When E2 binds to the apo receptor, H12 is displaced and H3 is locked in the 'out' conformation by the ligand binding within the LBP allosterically propagates stabilisation to tertiary structure. Note: for visualisation purposes the H1-H3 loop is hidden in the active E2-bound structures due to being highly dynamic; note that electron density for the loop is poorly defined in the crystal structure of both the apo and active E2-bound receptor.

**Table S1.** Crystallographic processing and refinement

| Statistic | apo *rf*ERα LBD (9MDV) |
|---|---|
| **Wavelength** | 0.9537 |
| **Resolution range** | 35.12 - 2.0 (2.072 - 2.0)[a] |
| **Space group** | P 1 21 1 |
| **Unit cell** | |
| **Dimensions (Å)** | 44.374 94.507 60.954 |
| **Angles (°)** | 90 96.857 90 |
| **Total reflections** | 220611 (20208) |
| **Unique reflections** | 33376 (3294) |
| **Multiplicity** | 6.6 (6.1) |
| **Completeness (%)** | 98.80 (98.62) |
| **Mean I/sigma(I)** | 8.47 (1.53) |
| **Wilson B-factor** | 27.26 |
| **R-merge** | 0.1471 (1.148) |
| **R-meas** | 0.1597 (1.256) |
| **R-pim** | 0.0613 (0.5001) |
| **CC1/2** | 0.997 (0.569) |
| **Reflections used in refinement** | 33350 (3293) |
| **Reflections used for R-free** | 1686 (187) |
| **R-work** | 0.2407 (0.3055) |
| **R-free** | 0.2867 (0.3448) |
| **Number of non-hydrogen atoms** | 4097 |
| **macromolecules** | 3772 |
| **ligands** | 24 |
| **solvent** | 301 |
| **Protein residues** | 477 |
| **RMS(bonds)** | 0.003 |
| **RMS(angles)** | 0.59 |
| **Ramachandran favored (%)** | 98.72 |
| **Ramachandran allowed (%)** | 1.28 |
| **Ramachandran outliers (%)** | 0 |
| **Rotamer outliers (%)** | 0.72 |
| **Clashscore** | 4.45 |
| **Average B-factor** | 29.35 |
| **macromolecules** | 29.05 |
| **ligands** | 36.82 |
| **solvent** | 32.54 |

[a]Statistics for the highest-resolution shell are shown in parenthesis

**Table S2.** SAXS experimental details

| **Data Collection** | **Specifications** |
|---|---|
| Beamline | BioSAXS, ANSTO |
| X-ray wavelength | 1 |
| Detector | Dectris Pilatus 2M |
| Exposure time (s) | 1 |
| q range (Å$^{-1}$) | 0.0098 to 7.2027 |
| Temperature (K) | 293 |

| **Size Exclusion Chromatography** | **Specifications** |
|---|---|
| Column | Cytiva S200 5/150 GL (3.2 mL) |
| Flow rate (mL/min) | 0.3 |
| Buffer composition | 20 mM Tris-HCl pH 8, 150 mM NaCl, 5% glycerol, 2 mM TCEP |
| Temperature (K) | 293 |

| **Data Analysis Software** | |
|---|---|
| Data reduction | PyFAI |
| Data processing | BioXTAS RAW |
| Model building | DAMMIF/N (ATSAS) |

**Table S3.** SAXS processing statistics

| Control | apo *rf*ERα | apo *h*ERα |
|---|---|---|
| **q_min** | 0.0112 | 0.0127 |
| **n_min** | 2 | 4 |
| **q_max** | 0.3 | 0.3 |
| **n_max** | 408 | 408 |
| **Dmax** | 92 | 74 |

| | apo *rf*ERα | | apo *h*ERα | |
|---|---|---|---|---|
| **Parameters** | **Rg** | **I/0** | **Rg** | **I/0** |
| **Guinier** | 25 | 8.29E-03 | 24.42 | 7.33E-03 |
| **Guinier Err.** | 0.0797 | 1.62E-05 | 0.0783 | 1.52E-05 |
| **P(r)** | 25.02 | 8.30E-03 | 24.08 | 7.30E-03 |
| **P(r) Err.** | 0.0956 | 1.55E-05 | 0.0457 | 1.15E-05 |
| **Total estimate** | 0.765 | | 0.9239 | |
| $\chi^2$ | 1.2055 | | 1.2235 | |
| **Alpha** | 50.067 | | 20.11 | |

**Table S4**. Markov model mean-first passage times between metastable states for the apo *rf*ERα LBD

| State | 1 | 2 | 3 | 4 | 5 | 6 | 7 | 8 |
|---|---|---|---|---|---|---|---|---|
| **1** | 0 | 1659.24 | 9840.67 | 1217.88 | 33213.59 | 11549.38 | 11151.93 | 11735.05 |
| **2** | 45615.63 | 0 | 24686.95 | 12697.05 | 94676.73 | 8982.98 | 8585.4 | 9168.65 |
| **3** | 29722.32 | 464.08 | 0 | 67.73 | 78581.3 | 10339.22 | 9941.77 | 10524.89 |
| **4** | 31737.28 | 362.68 | 10284.59 | 0 | 80442.66 | 10227.38 | 9829.92 | 10413.05 |
| **5** | 6422.38 | 2559.81 | 10960.71 | 2116.86 | 0 | 12440.58 | 12043.13 | 12626.25 |
| **6** | 144387.9 | 89016.64 | 123326.8 | 111593.6 | 193352.9 | 0 | 1451.64 | 148.45 |
| **7** | 142703.1 | 87331.69 | 121641.9 | 109908.8 | 191668 | 263.47 | 0 | 449.13 |
| **8** | 145312.4 | 89941.17 | 124251.3 | 112518.1 | 194277.4 | 701.9 | 2376.98 | 0 |

*Note: MFPTs are reported in nanoseconds (ns)*

**Table S5.** Markov model mean-first passage times between metastable states for the active E2-bound *rf*ERα LBD

| State | 1 | 2 | 3 | 4 | 5 | 6 | 7 | 8 | 9 | 10 | 11 |
|---|---|---|---|---|---|---|---|---|---|---|---|
| **1** | 0 | 43417.85 | 8350.18 | 2786.5 | 1232.84 | 9577.25 | 8296.26 | 12114.34 | 8887.11 | 15165.72 | 13462.29 |
| **2** | 23618.38 | 0 | 4748.36 | 1832.17 | 1581.35 | 10693.79 | 8636.22 | 12451.47 | 9224.09 | 15502.82 | 13799.4 |
| **3** | 20202.47 | 33365.83 | 0 | 1912.55 | 1400.61 | 10390.48 | 8459.51 | 12275.32 | 9047.97 | 15326.68 | 13623.26 |
| **4** | 33592.69 | 51335.59 | 20517.7 | 0 | 634.57 | 10191.52 | 7675.69 | 11488.82 | 8261.32 | 14540.14 | 12836.74 |
| **5** | 47396.09 | 68178 | 36677.02 | 14251.23 | 0 | 12049.4 | 6370.14 | 10161.47 | 6933.15 | 13212.4 | 11509.16 |
| **6** | 44117.77 | 65931.8 | 34262.91 | 12652.78 | 538.27 | 0 | 7198.94 | 11049.22 | 7823.77 | 14100.99 | 12397.4 |
| **7** | 81612.46 | 102418.3 | 70913.9 | 48518.88 | 23707.5 | 45017.7 | 0 | 5026.86 | 1909.38 | 8113.98 | 6395.84 |
| **8** | 100416.4 | 121218.5 | 89714.7 | 67317.32 | 42130.02 | 63877.65 | 17525.38 | 0 | 5233.48 | 2652.81 | 756.01 |
| **9** | 93970.44 | 114772.5 | 83268.75 | 60871.37 | 35688.1 | 57431.7 | 11063.27 | 2199.77 | 0 | 5274.97 | 3550.65 |
| **10** | 102279.2 | 123081.2 | 91577.48 | 69180.08 | 43987.9 | 65740.89 | 19423.46 | 2053.14 | 7167.23 | 0 | 460.42 |
| **11** | 101994 | 122796 | 91292.27 | 68894.88 | 43704.96 | 65455.46 | 19121.93 | 1585.2 | 6845.18 | 1732.32 | 0 |

*Note: MFPTs are reported in nanoseconds (ns)*

**Table S6.** Apo *rf*ERα LBD metastable state free energies

| State | Stationary dist. (π) | G/kT |
|---|---|---|
| **1** | 0.0014 | 6.57 |
| **2** | 0.0041 | 5.49 |
| **3** | 0.0048 | 5.34 |
| **4** | 0.0467 | 3.06 |
| **5** | 0.0252 | 3.66 |
| **6** | 0.0563 | 2.87 |
| **7** | 0.0682 | 2.68 |
| **8** | 0.7927 | 0.23 |

**Table S7.** +E2 rfERα LBD metastable state free energies

| State | Stationary dist. (π) | G/kT |
|---|---|---|
| **1** | 0.0088 | 4.725 |
| **2** | 0.0088 | 4.728 |
| **3** | 0.0198 | 3.912 |
| **4** | 0.0239 | 3.733 |
| **5** | 0.0343 | 3.371 |
| **6** | 0.0397 | 3.224 |
| **7** | 0.0411 | 3.1849 |
| **8** | 0.054 | 2.918 |
| **9** | 0.1291 | 2.0466 |
| **10** | 0.1755 | 1.7399 |
| **11** | 0.4645 | 0.7667 |
